# Comparative Analysis of Multiple Consensus Genomes of the Same Strain of Marek’s Disease Virus Reveals Intrastrain Variation

**DOI:** 10.1101/2023.09.04.556264

**Authors:** Alejandro Ortigas-Vasquez, Utsav Pandey, Daniel Renner, Chris Bowen, Susan J. Baigent, John Dunn, Hans Cheng, Yongxiu Yao, Andrew F. Read, Venugopal Nair, Dave A. Kennedy, Moriah L. Szpara

**Affiliations:** Departments of Biology, Center for Infectious Disease Dynamics and the Huck Institutes of the Life Sciences, Pennsylvania State University, University Park, Pennsylvania 16802, USA; Biochemistry and Molecular Biology, Center for Infectious Disease Dynamics and the Huck Institutes of the Life Sciences, Pennsylvania State University, University Park, Pennsylvania 16802, USA; Viral Oncogenesis Group, The Pirbright Institute, Woking, UK, GU24 0NF; United States Department of Agriculture, Agricultural Research Service, US National Poultry Research Center, Southeast Poultry Research Laboratory, Athens, Georgia 30605, USA; United States Department of Agriculture, Agricultural Research Service, US National Poultry Research Center, Avian Disease and Oncology Laboratory, East Lansing, Michigan 48823, USA

**Author notes:** **Corresponding Author** Dr. Moriah L. Szpara.

**Keywords:** Intrastrain, Diversity, Population, Genome, MDV, CVI988, Md5, Variation

## Abstract

Current strategies to understand the molecular basis of Marek’s disease virus (MDV) virulence primarily consist of cataloguing divergent nucleotides between strains with different phenotypes. However, each MDV strain is typically represented by a single consensus genome despite the confirmed existence of mixed viral populations. To assess the reliability of single-consensus interstrain genomic comparisons, we obtained two additional consensus genomes of vaccine strain CVI988 (Rispens) and two additional consensus genomes of the very virulent strain Md5 by sequencing viral stocks and cultured field isolates. In conjunction with the published genomes of CVI988 and Md5, this allowed us to perform 3-way comparisons between consensus genomes of the same strain. We found that consensus genomes of CVI988 can vary in as many as 236 positions involving 13 open reading frames (ORFs). In contrast, we found that Md5 genomes varied only in 11 positions involving a single ORF. Phylogenomic analyses showed all three Md5 consensus genomes clustered closely together, while also showing that CVI988_GenBank.BAC_ diverged from CVI988_Pirbright.lab_ and CVI988_USDA.PA.field_. Comparison of CVI988 consensus genomes revealed 19 SNPs in the unique regions of CVI988_GenBank.BAC_ that were not present in either CVI988_Pirbright.lab_ or CVI988_USDA.PA.field_. Finally, we evaluated the genomic heterogeneity of CVI988 and Md5 populations by identifying positions with >2% read support for alternative alleles in two ultra-deeply sequenced samples. We were able to confirm that both populations of CVI988 and Md5 were mixed, exhibiting a total of 29 and 27 high-confidence minor variant positions, respectively. We did not find any evidence of minor variants in the positions corresponding to the 19 SNPs in the unique regions of CVI988_GenBank.BAC_. Taken together, our findings confirm that consensus genomes of the same strain of MDV can vary and suggest that multiple consensus genomes per strain are needed in order to maximize the accuracy of interstrain genomic comparisons.

## Introduction

Marek’s disease virus (MDV) is an oncogenic, lymphoproliferative and neuropathic alphaherpesvirus affecting chickens (*Mardivirus gallidalpha 2*, Genus *Mardivirus*; Family *Herpesviridae*). It was first reported in 1907 by Hungarian veterinarian József Marek (1). Successful isolation of MDV in culture was only realized six decades later, closely followed by the development of the first vaccine, HPRS-16/Att (2–4). HPRS-16/Att was soon replaced by a better replicating, equally protective vaccine derived from strain FC126 of turkey herpesvirus (5). This vaccine, known as HVT, became the first widely used commercial vaccine for Marek’s disease (MD) (6). HVT was successful in reducing MD incidence in commercial flocks until the late 1970s, when reports of strains with increased virulence that were able to bypass vaccine-induced immunity started to emerge across the USA (7, 8). A bivalent vaccine consisting of HVT with MDV serotype 2 strain SB-1 was introduced in response. However, a similar shift in virulence occurred in just over a decade (9, 10). Outside the USA, a vaccine known as CVI988 had been in use since the mid 1970s (11). Developed in the Netherlands by a group of veterinary scientists under Dr. Bart Rispens, CVI988 offered the best protection against the emerging variants, and it quickly became the “gold standard” vaccine around the world (12). CVI988 is colloquially known today as the “Rispens” vaccine for MDV. Though CVI988 has mostly kept MD losses at bay for the past 30 years, there are now reports of strains potentially able to bypass CVI988-conferred immunity, and the most recent estimate places MDV-associated losses in the poultry industry at around US $1-2 billion per annum (13–16).

Comparative genomics has been a driving force in efforts to understand the molecular basis of MDV-induced virulence ever since the first viral genomes were sequenced over two decades ago (17, 18). The current strategy involves cataloging variable nucleotide positions between the consensus genomes of different MDV strains. Comparisons are made between strains grouped on the basis of their ability to break through vaccine protection, or “pathotype,” with the hope that this will help to identify the genes and polymorphisms associated with virulence (19, 20). There are defined pathotypes (i.e., mild/m, virulent/v, very virulent/vv, very virulent +/vv+), with CVI988 being classified as mild and Md5 as very virulent (21). However, despite MDV strains being known to exist as mixed populations, the degree to which the published consensus genomes represent the average for each strain has never been assessed (22–24). In addition, all available consensus genomes for MDV are low-coverage genomes that only account for the major variants (≥50%) present in the original viral population from which they were sequenced. The lack of information on minor variants (≤50%) can be problematic, due to previous reports suggesting that the diversity within a given viral population contributes to the overall pathotype of a strain (25–28). Although the extent of this contribution is still unknown, this nonetheless suggests that failing to account for minor variants can negatively impact our ability to correlate sequencing data to pathotype (21). Finally, many of the available reference genomes underwent cloning steps prior to sequencing. This includes the only available CVI988 consensus genome, which was sequenced from a bacterial artificial chromosome (BAC) copy of the viral genome, followed by plasmid-based cloning thereof (29). The Md5 consensus genome was obtained from plasmid-based cloning of viral DNA amplified in cell culture (18). The use of BAC and plasmid cloning approaches generates an artificially homogenous population of viral genomes, further moving away from the real-world biology of herpesvirus strains. In addition, by randomly selecting one viral haplotype from a population, these methods inherently reduce the chances of obtaining a representative consensus genome.

To address these issues, it is necessary to test the reliability of available MDV consensus genomes and assess the genomic heterogeneity of MDV populations using next-generation sequencing (NGS) approaches. Whole-genome resequencing (WGR) is a simple yet powerful approach that consists of comparing newly sequenced individuals against a reference (30). Here, we have adapted this strategy for viruses by sequencing multiple populations of the same strain and determining whether these yield a consistent consensus genome, under the assumption that large shifts in genomic variant frequencies will manifest as consensus-level differences. We independently sequenced two populations of the vaccine strain CVI988 and two populations of the very virulent prototype strain Md5, from a combination of viral stocks and cultured field isolates. The newly-sequenced genomes were compared with previously published consensus genomes using whole-genome pairwise alignments. Using this approach, we were able to detect differences between consensus genomes of each strain. Notably, we found 19 single-nucleotide polymorphisms (SNPs) in the unique genomic regions of CVI988_GenBank.BAC_ that were not present in CVI988_Pirbright.lab_ and CVI988_USDA.PA.field_. Next, we assessed the genomic heterogeneity of CVI988 and Md5 by analyzing ultra-deep Illumina sequencing data (>1,000× depth) and identified positions where at least 2% of the reads supported an alternative allele (31). This method enabled detection of minor allele frequency (MAF) variants and allowed us to confirm that the ultra-deep sequenced populations of CVI988 and Md5 were mixed. These findings highlight the need to sequence multiple consensus genomes of each MDV strain in order to ensure the accuracy of interstrain genomic comparisons and enhance our ability to link viral genotypes to virulence phenotypes.

## Methods

### Viral culture and DNA isolation of USDA CVI988 and Md5

The virulence phenotypes and targeted (amplicon-based) sequencing of USDA CVI988 and Md5 have been previously described by Dunn and colleagues (21). The sample we refer to as “CVI988_USDA.PA.field_” was collected as a field sample in 2010 in Pennsylvania (MDV 709B_v_2010_PA), and “Md5_USDA.lab_” was collected in Maryland in 1977 (MDV Md5_vv_1977_MD) (21). Chicken embryo fibroblasts (CEFs) were used to culture these virus stocks. Cultures were maintained in 1:1 mixture of Leibovitz’s L-15 and McCoy’s 5A (LM) media supplemented with fetal bovine serum (FBS), 200 U/ ml penicillin, 20 μg/ml streptomycin, and 2 μg/ml amphotericin B in a 37 °C, 5% CO_2_ incubator. Cells were plated with 4% FBS LM media and maintained in 1% FBS LM media. For storage as viral stocks, infected cells were suspended in freezing media composed of 10% DMSO, 45% FBS, and 45% LM media and kept in liquid nitrogen.

### DNA library preparation and sequencing of USDA CVI988 and Md5

DNA libraries for USDA CVI988 and Md5 were prepared for next generation sequencing using an Illumina Miseq platform as previously described (32–34). Briefly, genetic material, including viral nucleocapsid DNA, was quantified by Qubit (Invitrogen, CA) and by a virus specific qPCR. Total DNA was then acoustically sheared using a Covaris M220, with settings at 60 s duration, peak power 50, 10% duty cycle, at 4 °C. DNA was then processed through the Illumina TruSeq Nano DNA prep kit, using the manufactures recommendations, and further quantified by Qubit (Invitrogen, CA), Bioanalyzer (Agilent), and library specific qPCR (KAPA biosystems). Libraries were then diluted to 17pM per manufacturer’s recommendation and sequenced using version 3 paired-end (2×300bp length) chemistry.

### Viral culture and DNA isolation of Pirbright CVI988 and Md5

The CVI988 stock we refer to as “CVI988_Pirbright.lab_” was obtained from the Pirbright Institute and prepared from commercial CVI988 vaccine, sourced in the UK (Poulvac Marek CVI vaccine; Zoetis), following 2 passages in primary chicken embryo fibroblast cells (CEF). DNA was prepared from approximately 5×10^6^ cells using the DNeasy-96 kit (Qiagen, Hilden, Germany), according to the manufacturer’s instructions, and eluted in DNase-free water.

The Md5 stock we refer to as “Md5_Pirbright.lab_” was also obtained from the Pirbright Institute and derived from a 7^th^ duck embryo fibroblast passage stock kindly provided by Dr. A. M. Fadly (Avian Disease and Oncology Laboratory, USA). To amplify this stock, 5-day-old Rhode Island Red chickens were inoculated intra-abdominally with 1000 plaque forming units of virus. Lymphocytes isolated from spleens harvested at 14 days post-infection were cultured with CEF, the cell-associated virus was harvested at 7 days when cytopathic effect was clearly visible, and then further passed in CEF to produce virus stocks. DNA was prepared from approximately 5×10^6^ cells of the 9^th^ passage CEF stock using the DNeasy-96 kit, and eluted in DNase-free water.

### DNA library preparation and sequencing of Pirbright CVI988 and Md5

DNA isolated from each MDV culture was quantified for total DNA by Qubit (Invitrogen) and the amount of either Md5 DNA or CVI988 DNA was confirmed by a previously described qPCR assay (35). Samples were then acoustically sheared using Covaris and high-throughput, deep sequencing DNA library preparations were produced using the Kapa HyperPrep Library Kit (Roche) as per manufacturer’s protocol, with 14 PCR cycles. Then, samples were processed through a previously described oligo-enrichment procedure, where MDV-specific probes (36) (myBaits, Arbor Biosciences) were used with the myBaits Target Capture Kit (Arbor Biosciences) to enrich for MDV DNA (37). An additional PCR amplification (14 cycles) yielded libraries which were assessed for quality by Qubit, as well as a KAPA-specific qPCR, which allowed for accurate dilutions of libraries to 4 pM before sequencing. Finally, libraries were sequenced using v3 chemistry 300 bp × 300 bp paired-end sequencing on an Illumina MiSeq, as per the manufacturer’s instruction.

### Processing of sequencing reads and genome assembly

Viral reads were identified by aligning raw reads in FASTQ files against a local database of 97 MDV genomes using BLASTN with an E-value cutoff of 1×10^-2^ (see Supplementary Table 5 for strain names and accession numbers). The selected reads were then subjected to the quality control and preprocessing step (Step 1) of our published viral genome assembly (VirGA) workflow (38). These properly paired reads were then used for *de novo* assembly using metaSPAdes v3.14.0 (39). The resulting file containing the metaSPAdes scaffolds in FASTA format served as input for the remaining steps of VirGA (Steps 3-4), which include genome linearization, annotation and post-assembly quality assessments. For the reference-guided contig-ordering step, we used a trimmed version (TRL and TRS regions removed) of the published consensus genome of each strain (**Table 1**). New consensus genomes were also trimmed for downstream analyses. Tandem repeats were identified in trimmed consensus genomes using Tandem Repeats Finder (TRF) v4.09 (40). Manual verification and masking of tandem repeats was conducted using Geneious Prime v2023.0.4. To resolve *meq* at the consensus level, reads were mapped to modified reference genomes possessing different isoforms of *meq*. Local sequencing coverages were then used to determine the consensus-level isoform.

**Table 1:**
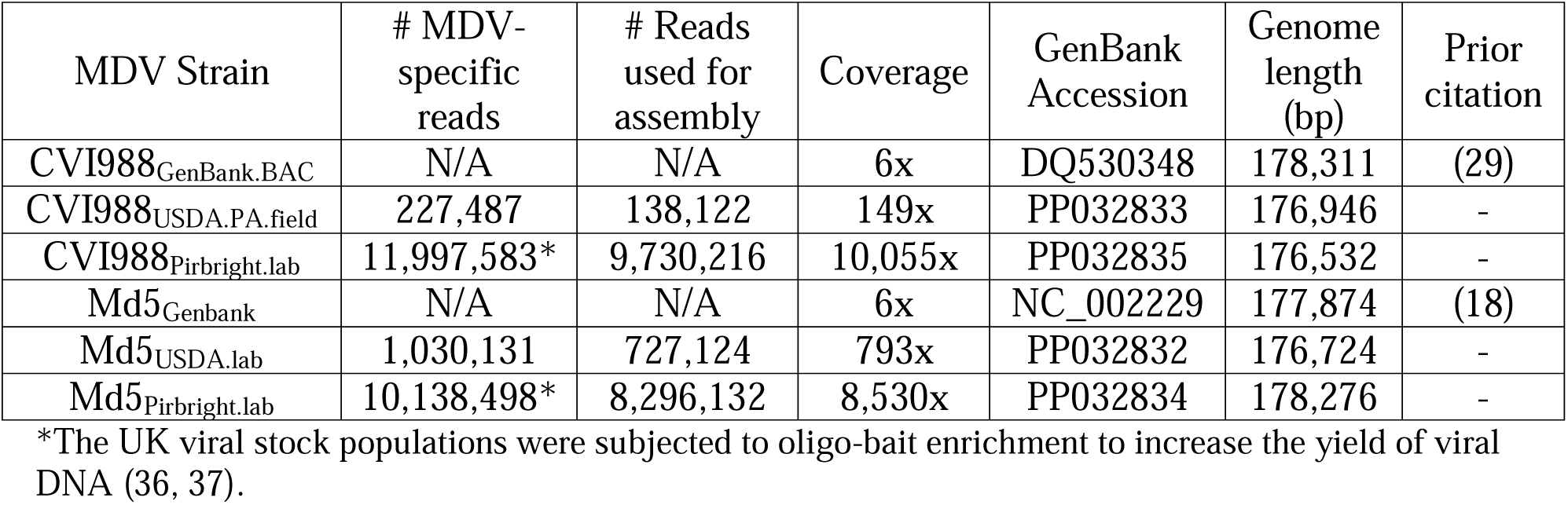
Sequencing statistics for new and previously published CVI988 and Md5 consensus genomes.

### Pairwise identity and genetic distance comparisons

Trimmed consensus genomes with masked tandem repeats were aligned using MAFFT v7.505 with default settings (41). The resulting alignment file in clustal format was imported into Geneious to visualize variable positions (SNPs and Indels) between the genomes. Published MDV consensus genomes sequenced from field samples or cultured stocks with PubMed Identifiers (n = 33) and all consensus genomes used for comparative analyses in the present study (n = 6) were aligned using MAFFT (see **Supplementary Table 5** for details). A neighbor-joining (NJ) tree was constructed from the multi-genome alignment file in clustal format using the Geneious Tree Builder tool with a Tamura-Nei genetic distance model and no outgroup.

### Identification of minor variant positions

A custom script was used to identify positions with at least 2% read support for an alternative allele – i.e., a minor variant – in genomes with >1,000x coverage. Positions in tandem repeats and homopolymers were excluded. All remaining minor variant positions were verified through manual inspection of BAM alignment files using the Integrative Genomics Viewer (IGV) v2.12.3 (42).

### Nucleotide sequence accession numbers

GenBank accession numbers for the four new and two prior viral genomes are listed in Table 1.

## Results

### Resequencing viral stocks and field isolates of CVI988 and Md5 using Illumina whole-genome sequencing

To enable intrastrain whole-genome comparisons, we first obtained two additional consensus genomes of both vaccine strain CVI988 (mild pathotype) and prototype very virulent strain Md5 (vv pathotype) (**Table 1**). Md5_USDA.lab_ and Md5_Pirbright.lab_ were sequenced from standard lab stocks (i.e., a virus population initiated from a prior viral stock and expanded in host cells).

CVI988_Pirbright.lab_ was likewise derived from lab stock. In contrast, CVI988_USDA.PA.field_ was isolated from a sample collected at a Pennsylvania farm, before isolation and expansion in host cells in the lab. Targeted, amplicon-based sequencing of portions of the Md5_USDA.lab_ (Md5_vv_1977_MD) and CVI988_USDA.PA.field_ (709B_v_2010_PA) genomes has been previously reported (21). All viral stocks were propagated similarly for viral DNA isolation and sequencing library preparation (see Methods for details). CVI988_Pirbright.lab_ and Md5_Pirbright.lab_ were additionally enriched using published methods for oligonucleotide-bait based enrichment in solution (36), which can enable a greater sequencing depth by enriching viral sequences from the overall milieu of DNA. Viral consensus genomes for each stock were determined using a previously published combination of *de novo* and reference-guided assembly methods (**Table 1**; see Methods for details).

The MDV genome has a double-stranded DNA (dsDNA) structure, with a full length of approximately 175-180 kb. The genomic layout is split into two large unique regions known as Unique Long (UL) and Unique Short (US), each flanked by inverted repeat sequences (IRL, IRS) and terminal repeat sequences (TRL, TRS) (**Figure 1A**, center). The α-type packaging sequences are located at the genomic termini and at the IRL/IRS junction. None of the new genomes contained any truncated or missing proteins. The average G+C content of all genomes was 44.0%. The average sequencing coverage of Illumina sequencing data exceeded 100× for all viral genomes and it exceeded 1,000× for the two genomes subjected to oligonucleotide-bait based enrichment (**Table 1**).

**Figure 1:**
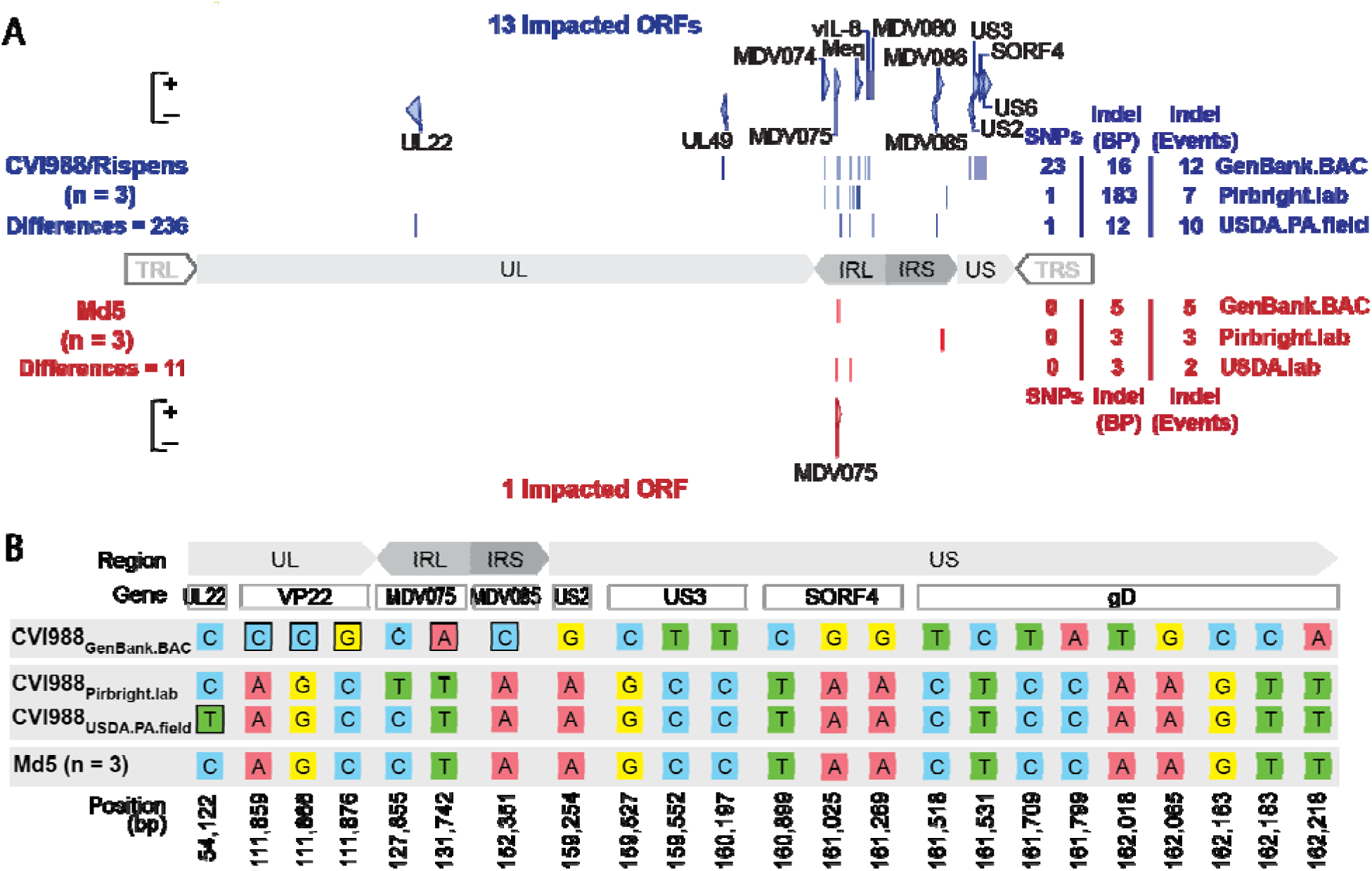
Comparative analysis of CVI988 and Md5 consensus genomes. **A)** A total of three trimmed consensus genomes (terminal repeat long (TRL) and short (TRS) removed) were obtained for each strain (CVI988 = Blue, Md5 = Red) and aligned using MAFFT. Consensus-level differences including SNPs and Indels are highlighted (dark lines) based on their relative locations in each respective genome (UL = Unique Long, IRL = Internal Repeat Long, IRS = Internal Repeat Short, US = Unique Short). The total number of differences between genomes of the same strain are indicated on the left, with a breakdown for each individual genome indicated on the right; these loci are all listed in detail in **Supplementary Table 1** and **Supplementary Table 2**. For each strain, ORFs with consensus-level differences in at least one of the three genomes belonging to that strain are highlighted and labeled (with alternative names when possible). The total number of ORFs with consensus-level differences is 13 for CVI988 and 1 for Md5. **B)** ORF-associated SNPs found between consensus genomes of CVI988. Genomic positions are indicated relative to CVI988_GenBank.BAC_, which accounted for 21 out of 23 ORF-associated SNPs. Completely unique SNPs that were not found in any of the other 38 MDV genomes included in the study are highlighted (black borders).

### Pairwise alignment of same-strain consensus genomes reveals greater intrastrain variation in CVI988 than Md5

For each strain, trimmed consensus genomes with masked tandem repeats were aligned and visualized. Consensus-level differences corresponding to SNPs, insertions and deletions (Indels) were identified and verified through manual inspection of sequence-read alignment files (i.e., BAM files). A total of 236 differences were detected in the three consensus genomes of vaccine strain CVI988, impacting a total of 13 ORFs. These include the genes UL22 / gH (MDV034), UL49 /VP22 (MDV062), MDV074, MDV075, Meq /RLORF7 (MDV076 / MDV005), vIL-8 / RLORF2 (MDV078 / MDV003), MDV080, MDV085, MDV086, US2 (MDV091), US3 (MDV092), SORF4 / S4 (MDV093) and US6 / gD (MDV094) (**Figure 1A, Supplementary Table 1**). Outside of regions associated with Indels and repetitive elements, CVI988_Pirbright.lab_ and CVI988_USDA.PA.field_ were found to be nearly identical (>99.99% identity), being distinguished only by a single synonymous SNP in UL22. In contrast, CVI988_GenBank.BAC_ possessed 23 unique SNPs and was the only CVI988 genome to exhibit variation in US. On the other hand, the three consensus genomes of very virulent strain Md5 showed a total of 11 differences, with only a single gene, MDV075, being impacted (**Supplementary Table 2**).

### The *meq* oncogene contributes significantly to overall intrastrain variation in consensus genomes of CVI988

Out of the 236 differences between CVI988 consensus genomes (**Supplementary Table 1**), 214 were found in IRL (n = 209 bp) and IRS (n = 5 bp). A large portion of these differences (n = 177 bp) corresponded to a 59 amino acid (AA) insertion in the *meq* oncogene, which was found in the CVI988_GenBank.BAC_ and CVI988_USDA.PA.field_ consensus genomes but absent in the CVI988_Pirbright.lab_ consensus genome (**Figure 2**). For *meq*, previous studies have suggested the existence of up to three isoforms: Meq, Long Meq (L-meq) and Short Meq (S-meq) (43–46).

**Figure 2:**
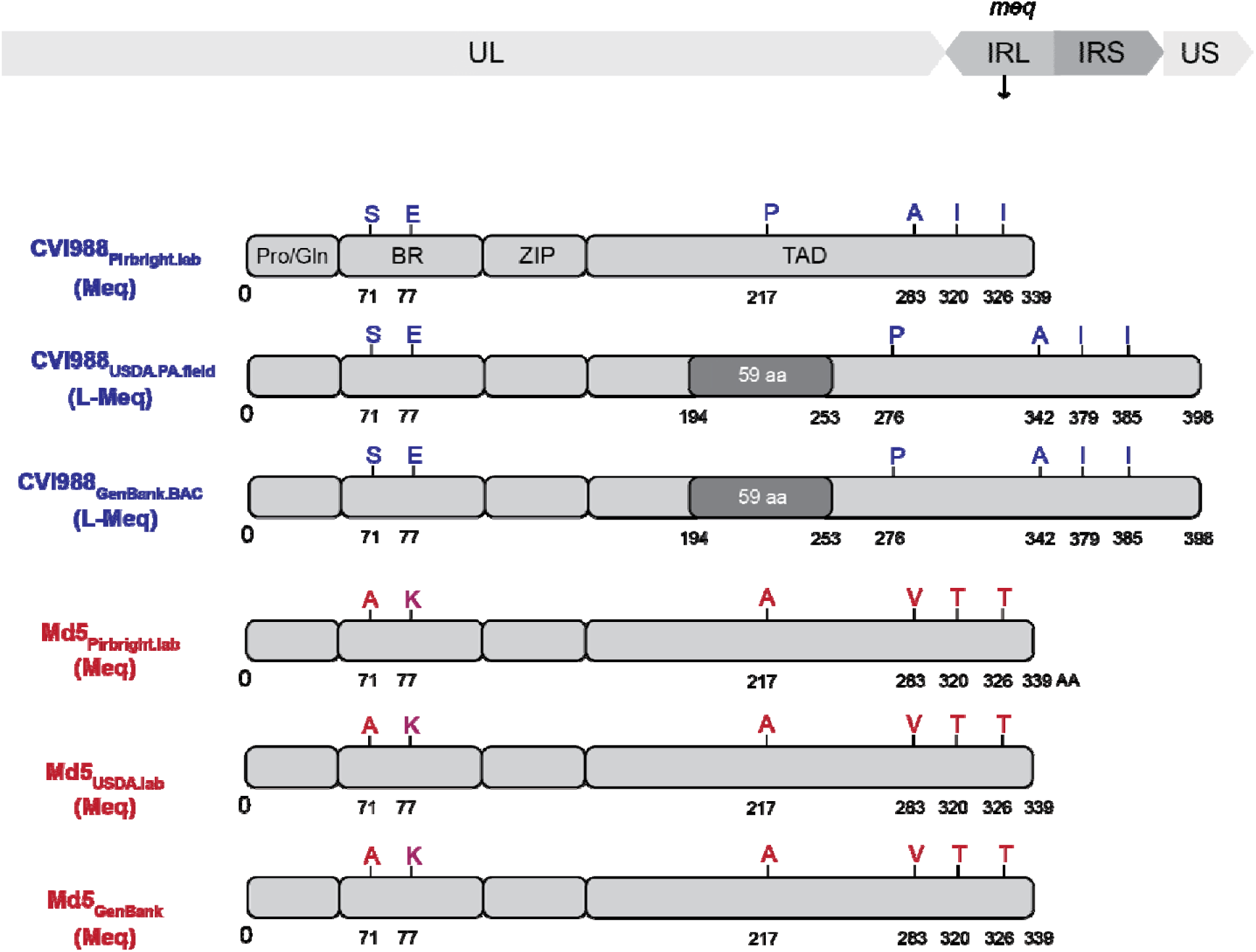
Comparison of the *meq* oncogene in Md5 and CVI988 consensus genomes. Schematic representation of IRL *meq* isoforms in CVI988 (blue) and Md5 (red) consensus genomes. The *meq* oncogene is comprised of the proline/glutamine (Pro/Gln) rich domain, the basic region (BR), the leucine zipper (ZIP) and the transactivation domain (TAD) at the C-terminal region. Differences in the AA sequence of the two strains are highlighted and labeled (A = Alanine, K = Lysine, S = Serine, E = Glutamic acid). The Meq protein is 339 AA in length. The L-Meq isoform contains a 59-amino acid insertion in the transactivation domain, resulting in a total length of 398 AA.

Meq is the standard isoform and is 339-AA in length. L-meq is a longer version of Meq that is characterized by a 59-AA insertion in the transactivation domain (TAD), which is often accompanied by a 3-bp CCA deletion immediately upstream from the insertion site. S-meq is a shorter version of Meq that exhibits a 41-AA deletion in the TAD. The CCA deletion that is typically associated with the presence of the L-meq isoform was confirmed to exist as a subpopulation of considerable size (∼ 40%) in the UK CVI988 population, but it did not appear to be linked to the presence of the 177-bp insertion (47). All three Md5 genomes exhibited the standard isoform of *meq* (**Figure 2**), again indicating less diversity when compared to CVI988.

### Increased intrastrain variation in structural repeat regions is associated with greater homopolymer instability

Outside of differences associated with *meq* isoforms, 28 of the 37 remaining differences found in IRS and IRL of CVI988 consensus genomes were associated with variation in homopolymer lengths (**Supplementary Table 1**). By contrast, out of the 22 differences found across the unique regions of CVI988 consensus genomes, 20 were ORF-associated SNPs, with an even distribution between synonymous (n = 10) and non-synonymous (n = 10). In Md5 consensus genomes, all 11 differences were found in either IRL (n = 6) or IRS (n = 5) and were all exclusively associated with variation in homopolymer lengths (**Supplementary Table 2**).

### Intrastrain variation in unique regions is associated with 19 SNPs in the published CVI988 consensus genome

Out of the 20 ORF-associated SNPs found in unique regions of CVI988 consensus genomes, 19 belonged to CVI988_GenBank.BAC_, including 3 SNPs in UL and 16 SNPs in US (**Figure 1B**). Phylogenomic analyses confirmed divergence of CVI988_GenBank.BAC_ from CVI988_Pirbright.lab_ and CVI988_USDA.PA.field_ (**Figure 3**). Additionally, none of the 3 SNPs found in UL of CVI988_GenBank.BAC_ were present in any of the other 38 MDV genomes included in this study. In contrast, all 16 SNPs in US of CVI988_GenBank.BAC_ were found in subsets across 6 different strains (EU-1, J-1, 814, GX0101, ATE2539 and Kgs-c1). However, no other strain apart from CVI988_GenBank.BAC_ individually possessed all 16 SNPs in the Unique Short (US) region. The nucleotides found across all 19 positions in CVI988_Pirbright.lab_ and CVI988_USDA.PA.field_ were also found in the three Md5 genomes (**Figure 1B**) and in 27 of the 33 additional MDV genomes used for phylogenomic analyses.

**Figure 3:**
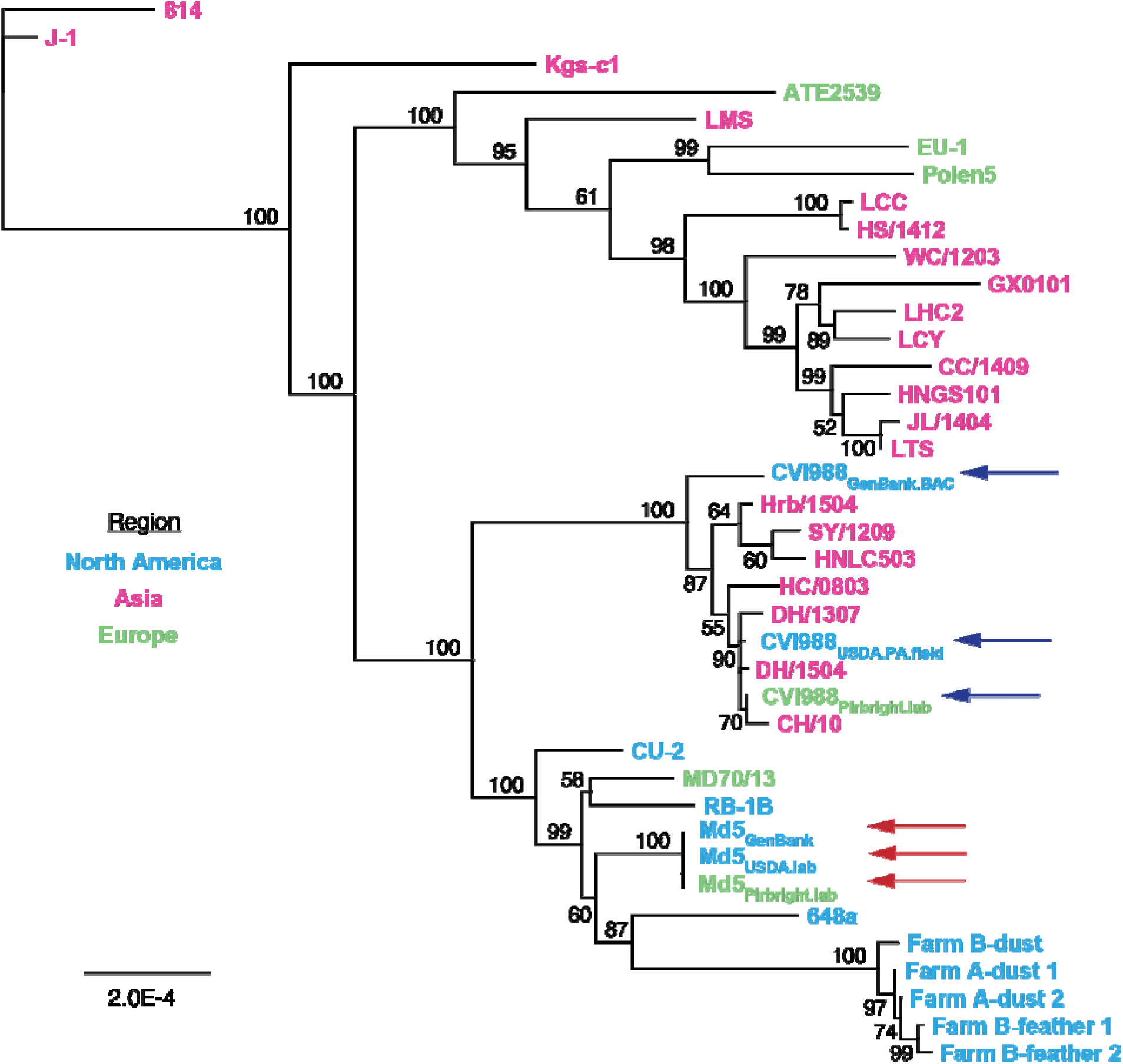
Neighbor-joining tree of previously published MDV strains and the new consensus genomes of CVI988 and Md5. Consensus genomes of CVI988, Md5 and 33 additional MDV strains from North America, Asia and Europe were aligned using MAFFT (**Supplementary Table 5**). The resulting MSA was used to infer a neighbor-joining tree using Geneious Prime v2023.0.4. CVI988 (blue arrows) and Md5 (red arrows) genomes used for comparative analyses are highlighted.

### Ultra-deep sequencing reveals presence of minor variants in CVI988 and Md5 populations

In addition to consensus-level comparisons, we performed minor variant analysis on genomes with a sequencing depth exceeding 1,000×. (**Figure 4**, see **Methods** for details). This included CVI988_Pirbright.lab_ (>10,000×) and Md5_Pirbright.lab_ (>8,500×; **Table 1**). A total of 29 minor variant positions were detected in the original CVI988_Pirbright.lab_ population across a combination of coding and regulatory regions. These minor variants impacted a total of 9 ORFs: MDV011, UL13 (MDV025), UL26 (MDV038), MDV069, MDV072, *meq* (MDV076), MDV077, ICP4 (MDV084) and US3 (MDV092) (**Supplementary Table 3).** Similarly, a total of 27 minor variants were detected in the UK Md5 population. These impacted a total of 10 ORFs: UL19 (MDV031), UL41 / VHS (MDV054), UL44 / gC (MDV057), UL48 / VP16 (MDV061), UL52 (MDV066), MDV069, MDV072, MDV078, MDV079 and MDV084 (**Supplementary Table 4**).

**Figure 4:**
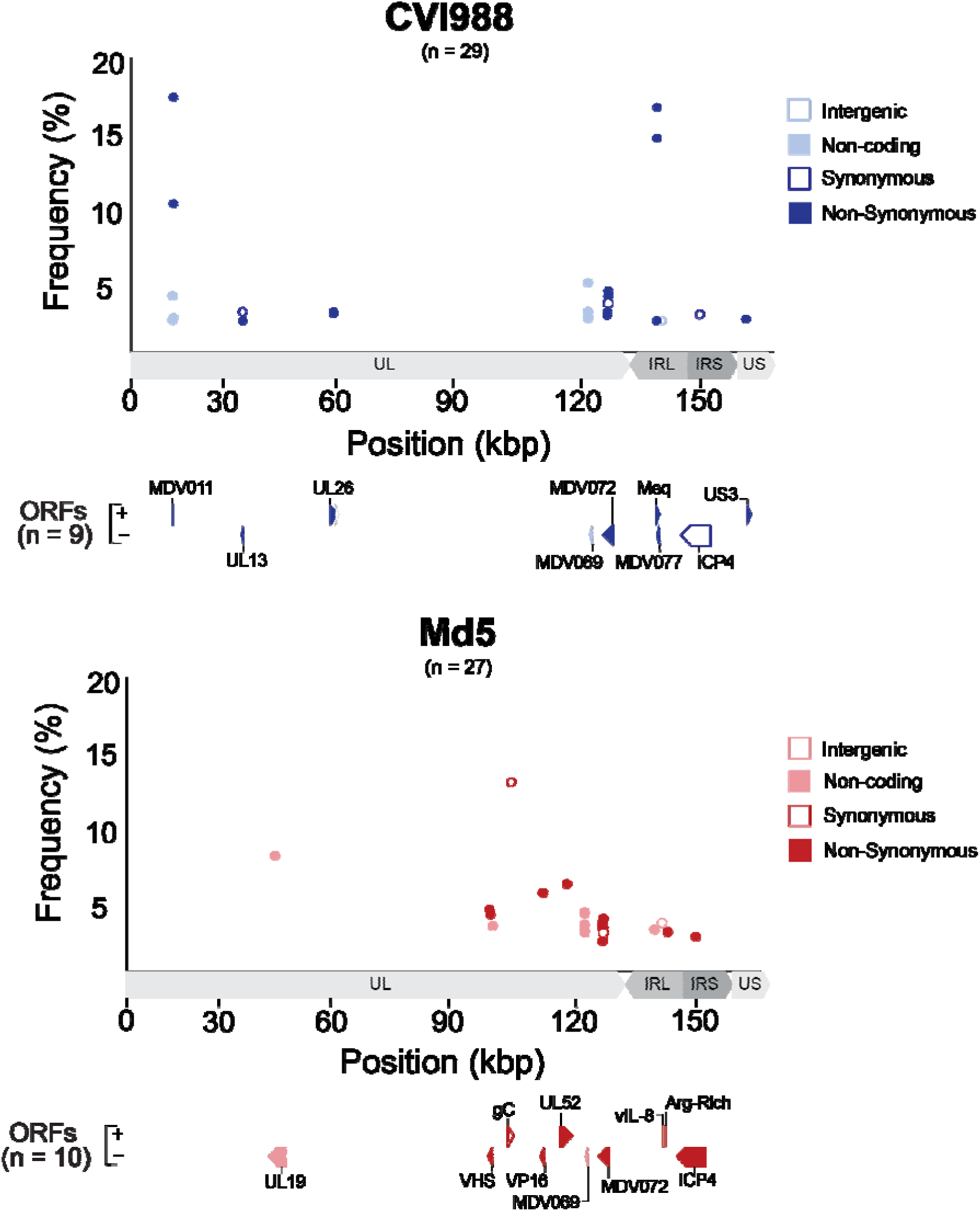
Minor variant distribution in Pirbright CVI988 and Md5 populations. For each strain (CVI988 = Blue, Md5 = Red), a single viral population was ultra-deep sequenced (>1,000×) to enable identification of minor variants (see **Table 1**). Positions outside of tandem repeats and homopolymers where the minor allele was present in ≥ 2% of the reads were identified as minor variants. Each minor variant was classified depending on its relative location to ORFs and its impact on the resulting amino-acid sequence (when applicable). ORFs associated with minor variants are labeled and highlighted beneath the graph using the same criteria.

Twelve of the 14 minor variant positions found across MDV069 and MDV072 were common to both strains. No minor variants were identified in the US region of the Md5_Pirbright.lab_ population. Additionally, we verified that no minor variant positions coincided with the 19 unique SNPs in CVI988_GenBank.BAC_ (**Figure 1B**).

## Discussion

The goal of comparative genomics is to provide a link between genotype and phenotype. The accuracy with which these two elements are represented is therefore directly proportional to our ability to make biologically relevant conclusions through the use of comparative genomic methods. DNA viruses have long been considered to be inherently stable, especially when compared to RNA viruses, which are now often described as mutant swarms, or “quasispecies” (48, 49). This had led to the assumption that viral consensus genomes, which represent the majority nucleotide at any given position, can accurately represent the genotype of MDV strains such as vaccine strain CVI988 and the prototype very virulent strain Md5. As demonstrated here and in prior studies, these and other herpesvirus populations can harbor minor variants both in culture and *in vivo* (50–53). Here, by sequencing additional viral stocks and field isolates of these strains, we have sought to evaluate the impact of conducting interstrain comparisons without accounting for the intrinsic genomic heterogeneity of MDV strain populations.

CVI988 is an attenuated and non-oncogenic vaccine strain that has long been known to exist as a mixed population due to undergoing multiple passages *in vitro*, in different laboratories and in multiple vaccine manufacturing companies. There have also been reports suggesting that the effectiveness of CVI988 vaccines can vary depending on the manufacturer, and that even different batches from the same manufacturer can show differences in effectiveness (54, 55). In contrast, Md5 is a virulent field strain of MDV that is currently used as the prototype very virulent (vv) strain in the USDA-ARS-Avian Disease and Oncology Laboratory (ADOL) pathotype classification system (56). As such, the fact that we found 235 differences between consensus genomes of CVI988 while only finding 11 differences between consensus genomes of Md5 is consistent with their history and real-world biology. Whether the larger number of variant loci in CVI988 reflects this strain’s attenuation through serial passage, or if it is a feature of this strain’s genetic background, remains to be determined. In addition, the fact that we were able to detect 29 minor variant positions in CVI988_Pirbright.lab_ and 27 minor variant positions in Md5_Pirbright.lab_ confirms that both strains exist as mixed viral populations. Over time, minor variant differences may expand during passage(s) in culture or during *in vivo* infections, leading to consensus level differences (50, 57). Minor variants may thus provide an explanation for the differences observed between consensus genomes of each strain.

Having access to the minor variant profiles of CVI988 and Md5 has the added advantage of providing insight into the potential effects of intrastrain variation in loci of high interest, such as pp38 (MDV073). Currently available qPCR assays to distinguish between CVI988 and virulent MDV strains are based on a single nucleotide polymorphism (SNP) located at position 320 of the pp38 gene (58). Our results show pp38 to be a highly conserved gene between different populations of CVI988 and Md5, with no minor variant positions coinciding with this gene. This not only confirms the robustness of qPCR assays based on the SNP #320 of pp38, but also establishes pp38 as a locus with high levels of within-strain conservation. Overall, our study provides a solid starting point for assessing intrastrain diversity in MDV strains. However, access to additional consensus genomes of CVI988 and Md5 will still be required in order to obtain the complete picture of MDV intrastrain diversity, as well as resequencing other virulent and attenuated MDV strains.

Among our most significant findings was the discovery of 19 SNPs present in the unique regions of CVI988_GenBank.BAC_ but absent in the two new CVI988 consensus genomes sequenced as part of this study. These 19 SNPs were also not present as minor variants in the CVI988_Pirbright.lab_ viral population, where these positions were sequenced at an average depth of >10,000×.

CVI988GenBank.BAC was sequenced by Spatz *et al*. in 2007 (29) from a BAC clone of CVI988 at 6× coverage (**Table 1**). This BAC clone was derived by Petherbridge *et al*. in 2003 from a commercial vaccine supplied by Fort Dodge Animal Health (59). As part of the cloning process, a BAC vector was inserted into US2 and the missing region was later reconstructed using PCR and wild-type CVI988 DNA. While explaining the origin of these 19 SNPs in CVI988_GenBank.BAC_ is beyond the scope of our current study, our results strongly suggest that consensus genomes of CVI988 stocks do not typically differ from virulent strains such as Md5, 648a and RB-1B in any of these positions.

The implications of these findings are multiple. Firstly, these 19 SNPs have been reported as differences between CVI988 and virulent strains like Md5, RB-1B, GA and Md11 by many prior studies, starting with the original publication by Spatz *et al.* (60–62). Secondly, the 16 unique SNPs in US of CVI988_GenBank.BAC_ impacting US2, US3, SORF4 and US6 / gD have been proposed as evidence of naturally occurring recombination between CVI988 and virulent strains (63). To date, a total of seven isolates from China have been described as natural recombinants of CVI988, largely on the basis of these 16 SNPs (64–66). The two new consensus genomes of CVI988 generated as part of this study share the same nucleotides in all 16 positions with these isolates, calling into question whether they are in fact natural recombinants of CVI988. An alternative explanation could be that these isolates are examples of freely circulating CVI988 that have reverted to virulence. In such a scenario, the remaining nucleotide differences between these isolates and our two new CVI988 consensus genomes could yield valuable information regarding loci associated with virulence. On the other hand, these isolates could also correspond to vaccine CVI988 reisolated from vaccinated birds. Either way, confirming the status of these seven isolates will require additional consensus genomes of CVI988, ideally from vaccine samples currently being distributed in China. Overall, these findings suggest that a heavy reliance on the published CVI988 consensus genome may have led to a historical overestimation of genomic divergence between CVI988 and virulent strains.

Finally, it is worth mentioning that current Illumina-based approaches do not provide insight into certain regions in MDV genomes where additional intrastrain variation is likely to be found. Specifically, tandem repeats have been suggested to be an important source of diversity in herpesviruses yet remain mostly inaccessible to NGS strategies both at the consensus and sub-consensus levels (52, 67). The use of long read technologies (e.g., Oxford Nanopore and Pacific Biosciences) could potentially help to overcome these limitations and make it possible to reliably assess intrastrain variation at tandem repeats and homopolymers (68, 69). In addition, the use of long reads could enhance our ability to link minor variant positions into proper haplotypes and help us to better understand the real-world biology of MDV strains.

In conclusion, by sequencing multiple viral stocks and field isolates of the vaccine strain CVI988 and the very virulent strain Md5, we have shown that consensus genomes of these strains can differ in multiple positions across all genomic regions. We have also established that these differences are most likely made possible by fluctuations in a mixed population of viral genomes, the existence of which we have demonstrated using ultra-deep Illumina sequencing. In addition, we have shown that not accounting for the intrinsic genomic heterogeneity of MDV strains may have led to a historical overestimation of divergence between CVI988 and virulent strains, providing an alternative explanation for sequencing data that was previously used to suggest naturally occurring recombination between the two. In doing so, we have demonstrated the value of working with multiple consensus genomes per strain and challenged prevailing notions regarding the genomic stability of DNA viruses.

## Acknowledgements

The authors thank members of the Szpara, Kennedy, and Nair labs for helpful feedback and discussion. We thank Matthew J. Jones and Andrew S. Bell for their contributions to early preparation of two samples for this study. This work was supported by NSF-NIH EEID award 1 R01 GM140459 (DK, MS, VN), the Biotechnology and Biological Sciences Research Council (BBSRC) grants BBS/E/I/00007039, BBS/E/PI/23NB0003, BB/CCG2250/1 and BB/V017748/1.

## CRediT (Contributor Roles Taxonomy; https://credit.niso.org/contributor-roles-defined/)

Conceptualization – AOV, MLS, DAK

Data curation – UP, DWR, AOV

Formal analysis – AOV

Funding acquisition – DAK, VN, MLS, AFR

Investigation – UP, CDB, SJB, YY, JD, HC

Methodology – UP, MJJ

Project administration – MLS, DAK

Resources – JD, HC, SJB, YY, VN, DWR, AFR

Software – UP, DWR, AOV

Supervision – MLS, DAK

Validation – CDB, DWR, AOV

Visualization – AOV, MLS, DAK

Writing – AOV

Writing – review & editing – AOV, MLS, DAK

## Supplementary Tables

**Supplementary Table 1:**
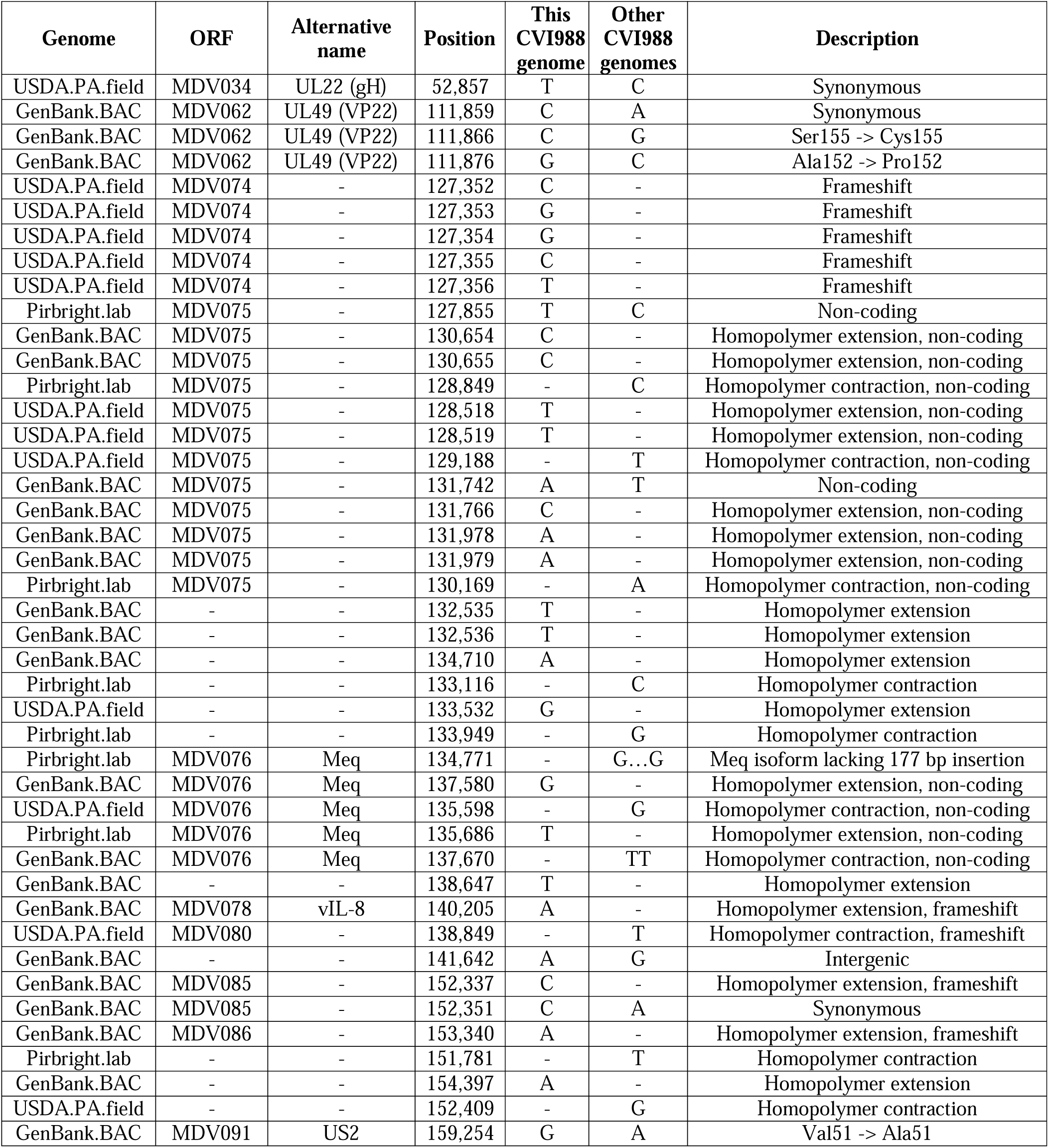
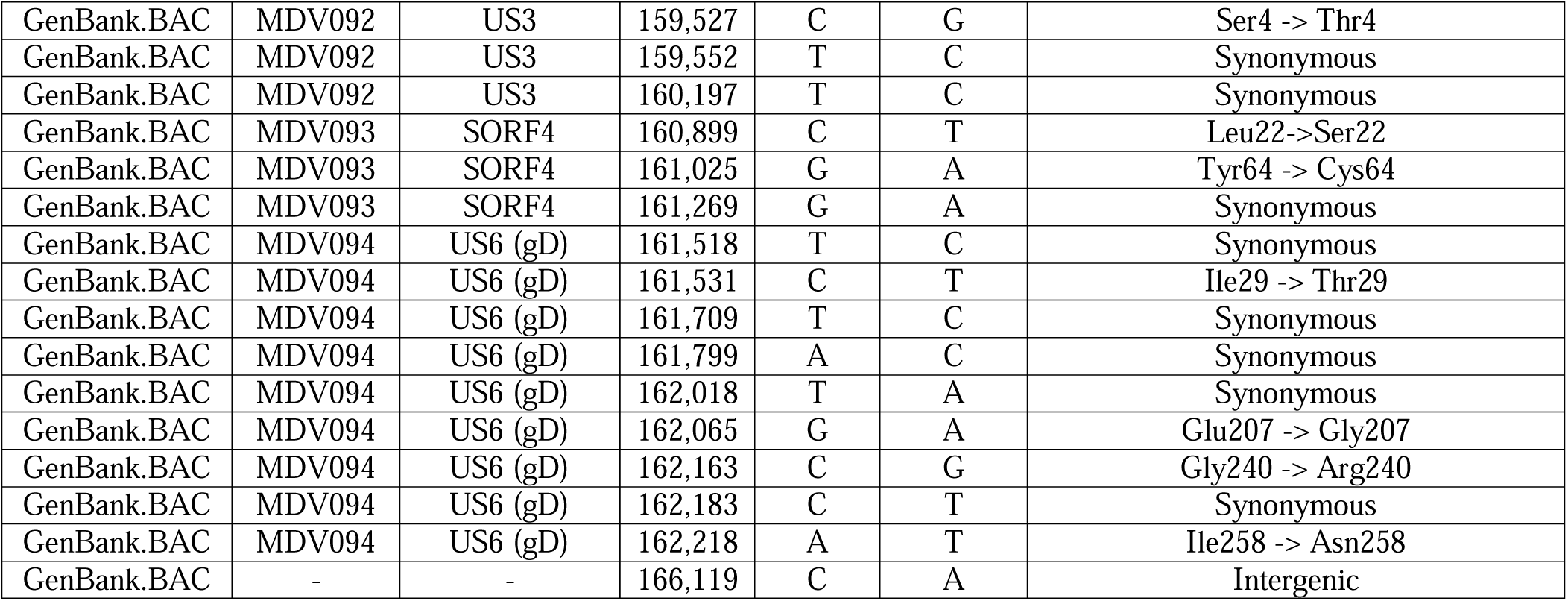
Variable positions in CVI988 consensus genomes.

**Supplementary Table 2:**
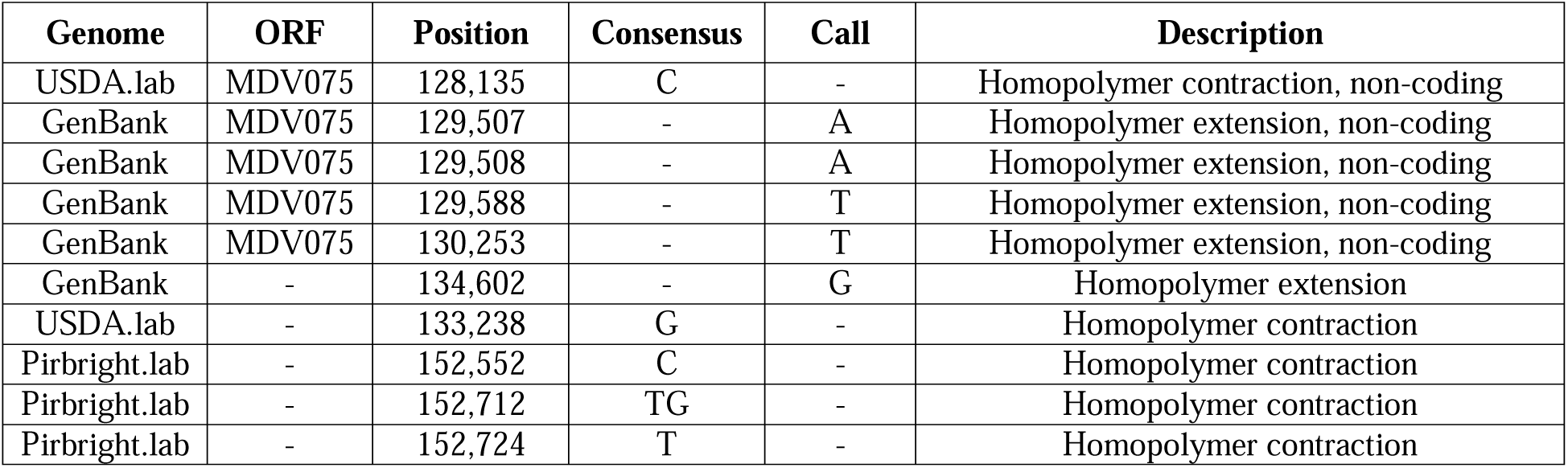
Variable positions in Md5 consensus genomes.

**Supplementary Table 3:** Minor variants in CVI988_Pirbright.lab_.

| Position in the genome | Major allele | Minor allele | Minor allele frequency | Major allele # forward reads | Major allele # reverse reads | Minor allele # forward reads | Minor allele # reverse reads | ORF | Type |
| --- | --- | --- | --- | --- | --- | --- | --- | --- | --- |
| 16732 | T | C | 4.65% | 5651 | 2646 | 275 | 130 | MDV011 | Upstream non-coding |
| 16735 | G | T | 2.22% | 5817 | 2636 | 126 | 66 | MDV011 | Upstream non-coding |
| 16770 | A | G | 3.21% | 5990 | 6705 | 199 | 222 | MDV011 | Upstream non-coding |
| 16809 | A | G | 17.30% | 4691 | 5465 | 919 | 1206 | MDV011 | Non-synonymous (Thr2 -> Ala2) |
| 16918 | A | G | 10.06% | 2353 | 2384 | 266 | 264 | MDV011 | Non-synonymous (Glu38->Gly38) |
| 36461 | A | T | 2.65% | 2845 | 3416 | 63 | 108 | MDV025 | Synonymous |
| 36463 | T | G | 2.09% | 2853 | 3418 | 46 | 88 | MDV025 | Non-synonymous (Thr265->Pro265) |
| 59022 | G | T | 2.55% | 4836 | 3741 | 72 | 153 | MDV038 | Non-synonymous (Ala239->Ser239) |
| 59040 | A | C | 2.70% | 4512 | 2768 | 61 | 141 | MDV038 | Non-synonymous (Thr245->Pro245) |
| 122353 | G | A | 2.11% | 6326 | 5563 | 138 | 119 | MDV069 | Upstream non-coding |
| 122362 | G | A | 2.26% | 6323 | 5494 | 151 | 123 | MDV069 | Upstream non-coding |
| 122405 | A | G | 2.34% | 5802 | 5540 | 145 | 128 | MDV069 | Upstream non-coding |
| 122419 | T | C | 2.66% | 3553 | 5549 | 127 | 122 | MDV069 | Upstream non-coding |
| 122421 | T | G | 2.66% | 3552 | 5519 | 122 | 126 | MDV069 | Upstream non-coding |
| 122848 | A | G | 2.43% | 4877 | 5946 | 116 | 154 | MDV072 | Non-synonymous (Leu875->Pro875) |
| 122857 | A | G | 2.66% | 4802 | 5647 | 123 | 163 | MDV072 | Non-synonymous (Val872->Ala872) |
| 122900 | G | A | 3.76% | 3676 | 4473 | 141 | 178 | MDV072 | Non-synonymous (His858->Tyr858) |
| 122914 | C | T | 3.25% | 3332 | 4093 | 81 | 169 | MDV072 | Non-synonymous (Cyst853->Tyr853) |
| 122916 | G | T | 3.27% | 3270 | 4023 | 80 | 167 | MDV072 | Synonymous |
| 122977 | T | G | 4.08% | 2886 | 2890 | 84 | 162 | MDV072 | Non-synonymous (His832->Pro832) |
| 123012 | C | T | 3.28% | 2684 | 2155 | 51 | 113 | MDV072 | Non-synonymous (Trp820->STOP) |
| 134751 | A | G | 14.59% | 25181 | 19947 | 5447 | 2316 | MDV076 | Non-synonymous (Thr180->Ala180) |
| 134759 | A | G | 16.71% | 23815 | 19819 | 6391 | 2431 | MDV077 | Non-synonymous (Phe111->Leu111) |
| 134784 | C | - | 58.08% | 29714 | 19086 | 14302 | 14301 | MDV076 | Deletion |
| 134785 | C | - | 57.02% | 30172 | 19826 | 14287 | 14286 | MDV076 | Deletion |
| 134786 | A | - | 57.12% | 29125 | 19029 | 14288 | 14287 | MDV076 | Deletion |
| 137914 | T | C | 2.20% | 3776 | 3761 | 86 | 84 | - | Intergenic |
| 144482 | T | C | 2.60% | 11669 | 13033 | 441 | 219 | MDV084 | Synonymous |
| 157396 | A | G | 2.16% | 6886 | 5749 | 157 | 123 | MDV092 | Non-synonymous (Glu54->Gly54) |

**Supplementary Table 4:** Minor variants in Md5_Pirbright.lab_.

| Position in the genome | Major allele | Minor allele | Minor allele frequency | Major allele # forward reads | Major allele # reverse reads | Minor allele # forward reads | Minor allele # reverse reads | ORF | Type |
| --- | --- | --- | --- | --- | --- | --- | --- | --- | --- |
| 49848 | A | G | 7.89% | 3400 | 3374 | 278 | 303 | MDV031 | Non-coding |
| 97742 | T | G | 4.15% | 2275 | 1949 | 107 | 76 | MDV054 | Non-synonymous (His136->Pro136) |
| 97829 | G | T | 3.82% | 2575 | 1105 | 106 | 40 | MDV054 | Non-synonymous (Thr107->Asn107) |
| 98543 | T | C | 3.09% | 1091 | 3673 | 33 | 119 | MDV054 | Non-coding |
| 102545 | A | G | 12.92% | 3903 | 2728 | 590 | 397 | MDV057 | Synonymous |
| 110386 | C | T | 5.32% | 4509 | 2255 | 259 | 121 | MDV061 | Non-synonymous (Asp35->Asn35) |
| 114480 | G | A | 5.96% | 2422 | 4690 | 229 | 222 | MDV066 | Non-synonymous (Val192->Ile192) |
| 122515 | G | A | 2.61% | 4183 | 3701 | 53 | 158 | MDV069 | Non-coding |
| 122524 | G | A | 2.80% | 4122 | 3688 | 65 | 160 | MDV069 | Non-coding |
| 122567 | A | G | 3.17% | 3769 | 3694 | 68 | 177 | MDV069 | Non-coding |
| 122581 | T | C | 3.90% | 2128 | 3659 | 63 | 172 | MDV069 | Non-coding |
| 122583 | T | G | 4.03% | 2122 | 3624 | 63 | 179 | MDV069 | Non-coding |
| 122645 | G | T | 3.05% | 2982 | 3659 | 51 | 158 | MDV069 | Non-coding |
| 122977 | G | A | 2.01% | 3501 | 4391 | 92 | 70 | MDV072 | Non-synonymous (Ala886->Val886) |
| 123010 | A | G | 2.97% | 3503 | 4076 | 142 | 90 | MDV072 | Non-synonymous (Leu875->Pro875) |
| 123019 | A | G | 3.29% | 3495 | 3995 | 158 | 97 | MDV072 | Non-synonymous (Val872->Ala872) |
| 123062 | G | A | 3.57% | 2986 | 3374 | 151 | 85 | MDV072 | Non-synonymous (His858->Tyr858) |
| 123076 | C | T | 2.55% | 2852 | 3217 | 78 | 81 | MDV072 | Non-synonymous (Cyst853->Tyr853) |
| 123078 | G | T | 2.57% | 2803 | 3185 | 76 | 82 | MDV072 | Synonymous |
| 123139 | T | G | 3.58% | 2753 | 2417 | 110 | 82 | MDV072 | Non-synonymous (His832->Pro832) |
| 123174 | C | T | 2.87% | 2726 | 2034 | 90 | 51 | MDV072 | Non-synonymous (Trp820->STOP) |
| 132447 | C | T | 2.68% | 6309 | 4595 | 161 | 139 | - | Intergenic |
| 137705 | G | A | 3.12% | 5261 | 3764 | 168 | 123 | MDV078 | Synonymous |
| 138785 | G | C | 2.55% | 158 | 301 | 4 | 8 | MDV079 | Non-synonymous (Ala107->Pro107) |
| 145343 | T | G | 2.25% | 6893 | 3297 | 137 | 98 | MDV084 | Non-synonymous (Glu1725->Ala1725) |
| 147827 | G | A | 21.44% | 3490 | 2598 | 963 | 699 | MDV084 | Non-synonymous (Ala897->Val897) |
| 150329 | T | C | 31.19% | 7233 | 6650 | 3294 | 3002 | MDV084 | Non-synonymous (Gln63->Arg63) |

**Supplementary Table 5:**
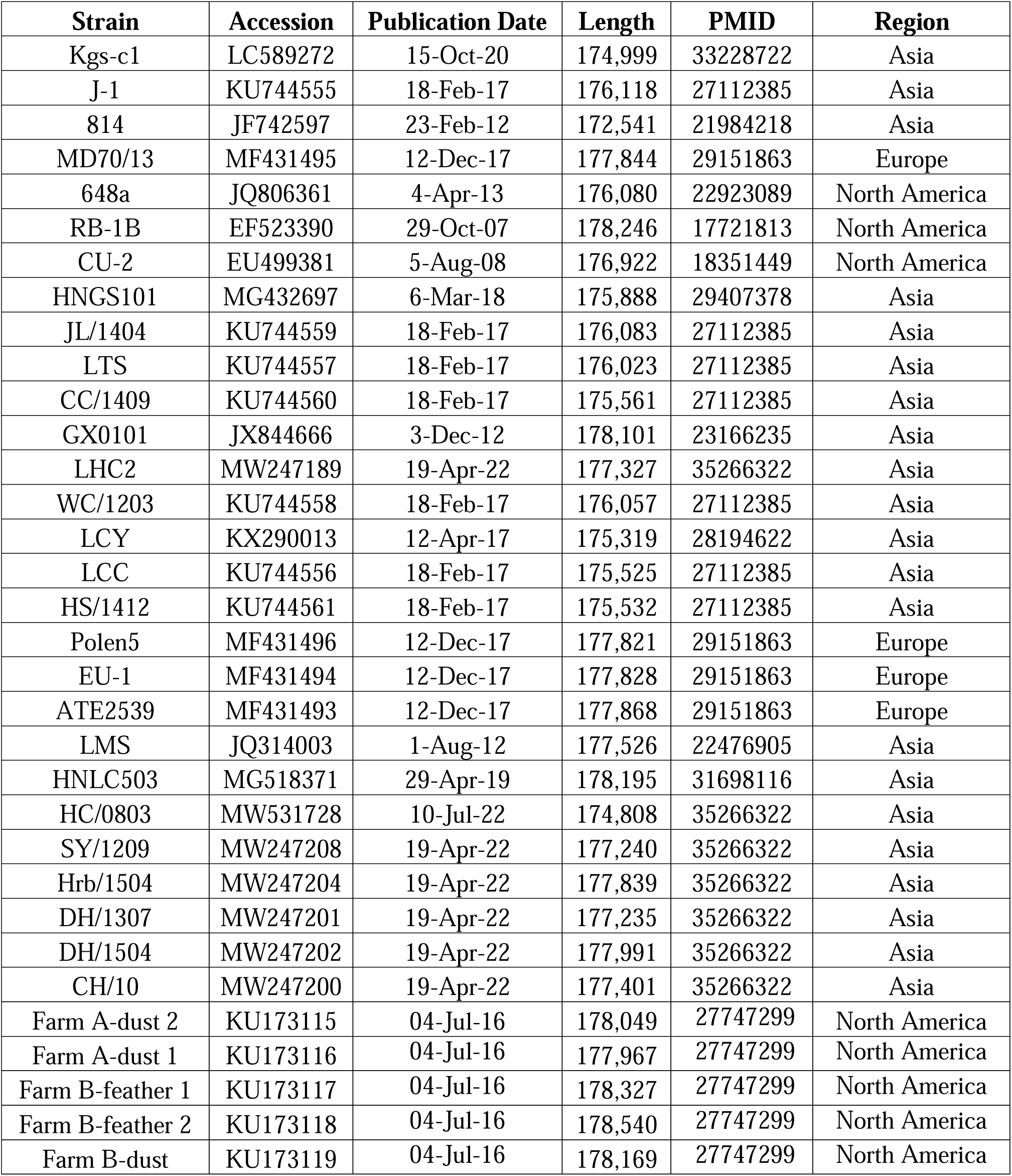
Published reference genomes for additional 33 strains used to construct NJ tree.

